# Direct regulation of cell cycle regulatory gene expression by NtrX to promote *Sinorhizobium meliloti* cell division

**DOI:** 10.1101/2020.11.06.341933

**Authors:** Shenghui Xing, Fang An, Leqi Huang, Xinwei Yang, Shuang Zeng, Ningning Li, Lanya Zhang, Wenjia Zheng, Khadidja Ouenzar, Junhui Yan, Liangliang Yu, Li Luo

## Abstract

Cell division of the alfalfa symbiont, *Sinorhizobium meliloti*, is regulated by a signaling network centered on CtrA. The gene expression of regulatory proteins in the network can be regulated by nutrient signaling systems. In this study, we found that NtrX, one of the regulators of nitrogen metabolic response, can directly regulate the expression of several regulatory genes in the CtrA signaling network. Three groups of *S. meliloti ntrX* mutants showed similar cell division defects, such as slow growth, abnormal morphology of some cells and delayed DNA synthesis. Quantitative RT-PCR assays indicated that in these mutants, the transcription of genes such as *ctrA* and *gcrA* was up-regulated, while the transcription of genes such as *dnaA* and *ftsZ1* was down-regulated. Western blotting showed that the CtrA and GcrA proteins were apparently increased in the mutants. The 53^rd^ aspartate conserved in NtrX homologs can be phosphorylated *in vitro* and *in vivo*. The phosphorylated NtrX protein can bind directly to the promoter regions of *ctrA, gcrA, dnaA* and *ftsZ1* by recognizing the characteristic sequence CAAN_1-5_TTG. Therefore, phosphorylation of NtrX is essential for cell cycle regulation of *S. meliloti*. We expressed the NtrX protein carrying a phosphorylation site substitution in *Agrobacterium tumefaciens* and found that the expressed strains had different growth phenotypes, suggesting that NtrX also regulates cell division in this bacterial species. Our findings reveal that NtrX acts as a transcriptional regulator that positively affects bacterial cell division, associated with nitrogen metabolism.

**IMPORTANCE:** *Sinorhizobium meliloti* infects the host legume alfalfa and induces the formation of nitrogen-fixing nodules. The proliferation of rhizobia in plant tissues and cells is strictly controlled in the early stage of the interaction between symbiotic partners. However, the control mechanism is not very clear. Cell division of *S. meliloti* in the free-living state is regulated by the CtrA signaling network, but the molecular mechanisms by which the CtrA system is associated with environmental nutrient signals (e.g., ammonia nitrogen) need to be further explored. This study demonstrates that NtrX, a regulator of nitrogen metabolism, required for symbiotic nodulation and nitrogen fixation by *S. meliloti* 1021, can act as a transcriptional regulator of the CtrA signaling system. It may link nitrogen signaling to cell cycle regulation in *Rhizobium* species.

---

*Caulobacter crescentus* is a model strain of α-proteobacteria in molecular cell biology (1). It takes advantage of one cell division to produce two cells with different shapes and sizes (2). In recent years, a complex cell cycle regulatory network has been revealed in this species. This network consists of multiple histidine kinases such as CckA, DivL, DivJ, and PleC, a histidine phosphotransfer protein ChpT, response regulators DivK and CpdR, and transcription regulators CtrA, GcrA, DnaA, SciP, and MucR (1–6). Although this network has been reported to possibly mediate nutritional signals for regulating bacterial growth and proliferation, the exact molecular mechanism is currently unclear.

*Sinorhizobium meliloti* is a model strain of rhizobia that can infect the host plant and form nitrogen-fixing nodules. During symbiosis, the cell division of *S. meliloti* on the surface of host plant alfalfa roots, at the front ends of extended infection threads and in the infection zones of root nodules, is stringently controlled (7), but how cell division is regulated is not very clear. Although NCR (Nodule Cysteine Rich) peptides secreted by host plants are known to induce terminal differentiation of bacteroids in host plant cells (8–11), several legumes do not produce NCR peptides. Therefore, there may be other mechanisms that control cell division of symbiotic rhizobia in host plants. Since *S. meliloti* and *C. crescentus* belong to α-proteobacteria, based on the research results of *C. crescentus*, with the aid of DNA sequence homology analysis, many cell cycle regulatory genes such as *ctrA, ccrM, cpdR1, divJ, divK, gcrA* and *pleC*, have been identified in *S. meliloti* (12–14). In addition, the hybrid histidine kinase CbrA is linked to the CtrA regulatory system, which is an important regulator of cell division in *S. meliloti* (15, 16). However, it is still unclear whether these regulatory proteins conduct environmental nutrition signals (e.g., ammonia nitrogen) and whether they play a regulatory role in the symbiotic process.

The NtrY/NtrX two-component system, first discovered in *Azorhizobium caulinodans*, regulates nitrogen metabolism under free-living conditions and affects nodulation and nitrogen fixation in the host plant *Sesbania rostrata* (17). Subsequently, *ntrY/ntrX* homologous genes were found in *Rhizobium tropici* to regulate nitrogen metabolism and symbiotic nodulation (18). NtrY/NtrX homologs regulate nitrate uptake in *Azospirillum brasilense* and *Herbaspirillum seropedicae* (19, 20), and this regulatory system has been found to simultaneously control nitrogen metabolism and cellular redox homeostasis in *Rhodobacter capsulatus* (21). Moreover, NtrX is involved in the regulation of cell envelope formation in *R. sphaeroides*(22). In *Brucella abortus*, the histidine kinase NtrY participates in micro-oxygen signaling and nitrogen respiration (23), while the response regulator NtrX controls the expression of respiratory enzymes in *Neisseria gonorrhoeae* (24). Interestingly, the NtrY/NtrX system regulates cell proliferation, amino acid metabolism and CtrA degradation in *Ehrlichia chaffeensis* (25). Finally, NtrX is required for the survival of *C. crescentus* cells and its expression is induced by low pH (26). These findings show that NtrY/NtrX appears to be a regulatory system for nitrogen metabolism, which may be involved in the regulation of cell division.

NtrX is a NtrC family response regulator protein, which consists of a receiver (REC) domain and a DNA-binding domain (27, 28). X-ray crystal diffraction results indicate that the NtrX protein of *B. abortus* can form a dimer; the REC domain is composed of 5 α-helices and 5 β-sheets; the DNA-binding domain contains an HTH motif, which includes 4 α-helices. The three-dimensional structure of the C-terminus has not been resolved (28). *In vitro* experiments demonstrated that the NtrX protein of *B. abortus* can recognize and bind to the palindromic DNA sequence (**CAAN**_3-5_**TTG**) in the *ntrY* promoter region via the HTH motif to regulate gene transcription (27, 28). In *S. meliloti* 1021, our previous work found that NtrX protein can regulate bacterial growth and proliferation, flagellar synthesis and motility, succinoglycan production, and symbiotic nodulation and nitrogen fixation with the host plant alfalfa (29, 30). In the present study, we investigated the control mechanism by which NtrX regulates *S. meliloti* cell division at the transcriptional level.

## RESULTS

### Defects of cell division resulting from *ntrX* mutation in *S. meliloti*

We previously constructed a plasmid insertion mutant of the *ntrX* gene in the background of *S. meliloti* 1021, SmLL1 (29). This mutant grew slowly even in the rich LB/MC medium compared to wild-type Sm1021 (29). According to the determined growth curve, the doubling time of bacterial cell proliferation was calculated to be 180 min for SmLL1 compared to 160 min for the wild-type strain as reported (34). Microscopic observation revealed that 5% to 10% of SmLL1 cells grown in the LB/MC broth up to the logarithmic phase exhibited morphological abnormalities (such as cell elongation, Y-shaped or V-shaped), whereas Sm1021 had almost no abnormally shaped cells (Fig. 1A). To determine whether the appearance of abnormally shaped cells of the SmLL strain is associated with the synthesis and segregation of genomic DNA, we synchronized the *S. meliloti* cells, inoculated them in LB/MC broth, grew them for 90 min and 180 min, respectively, and then harvested the cells for flow cytometric analysis. The results showed that Sm1021 synchronized cells were haploid containing 1C genomic DNA, while cells grown for 90 min were diploid containing 2C genomic DNA; cells grown for 180 min were both haploid and diploid (Fig. 1B). Cells of the SmLL mutant after synchronization were mainly haploid, and cells grown for 90 min were mainly diploid, but the cells grown for 180 min were still mainly diploid (Fig. 1B), indicating a decelerated of replication and segregation of their genomic DNA as compared to the wild type. These observations indicate that the SmLL mutant has cell division defects.

**Fig. 1.**
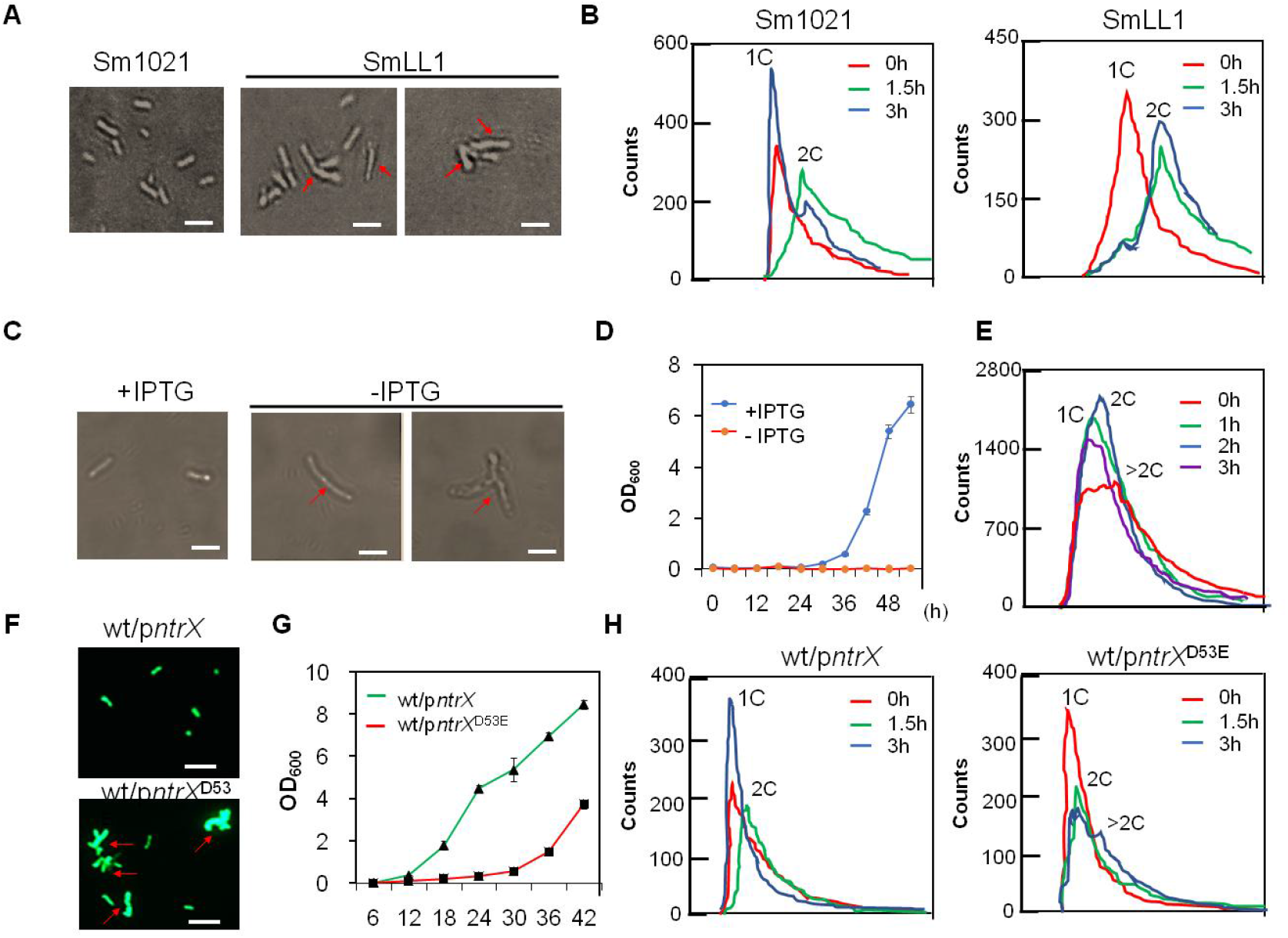
Cell division defects of *S. meliloti ntrX* mutants in LB/MC medium. (**A, C, F**) Cell shapes and sizes of *ntrX* mutants under a light or fluorescence microscope. Red arrows, abnormal cells; bars, 2μm in **A** and **C**; Bar, 100 μm in **F.** (**B, E, H**) Genomic DNA content of *ntrX* mutants was determined by flow cytometry. 1C, haploid; 2C, diploid; >2C, polyploid. (**D, G**) Growth curves of the *ntrX* depletion strain and Sm1021/p*ntrX*^D53E^. Error bars, ±SD. 1 mM IPTG was added to the LB/MC broth. SmLL1, *ntrX::pK18mobsacB;* wt, Sm1021.

It may be fatal due to the deletion of *ntrX*, and we have not yet successfully screened the deletion mutant of this gene in *S. meliloti* 1021, so a depletion strain (where the *ntrX* gene on the genome has been deleted and the NtrX protein in the cell is expressed from an IPTG-inducible plasmid) was constructed to verify the above results. Optical microscopic observation showed that more than 30% of the cells of the depleted strain in LB/MC broth without IPTG showed abnormal shapes (such as elongation and T-shaped), while in LB/MC broth with IPTG, almost no abnormal cells were observed (Fig. 1C). The depletion strain barely proliferated in LB/MC broth without IPTG (Figure 1D), indicating that *ntrX* gene expression is required for the cell division of *S. meliloti*. Flow cytometric analysis showed that in addition to haploid and diploid cells, polyploid cells were detected in the depletion strain in LB/MC broth without IPTG (Fig.1E). After the addition of IPTG, polyploid cells were almost undetectable (Fig. 1E), indicating that genomic DNA replication and segregation of *S. meliloti* requires the expression of the *ntrX* gene.

NtrX, as a regulator of nitrogen metabolism, is composed of a REC domain and a DNA-binding domain (28). The research results from *C. crescentus* and *B. abortus* showed that the D53 residue on the REC domain is the phosphorylation site of the protein (26, 28). We found that the REC domain of the NtrX protein from *S. meliloti* also contains a conserved D53 residue. If the NtrX protein is indeed involved in the regulation of *S. meliloti* cell division, as described above, then the mutation of the D53 residue would affect its regulatory function. To test this hypothesis, we began constructing amino acid substitution mutations (the codon of D was replaced by A, N or E, respectively) in the *ntrX* gene on the genome of *S. meliloti* 1021, but were unable to screen the mutants. As a result, we cloned the mutation gene into the expression vector pSRK-Gm (31) and then introduced the recombinant plasmid into Sm1021. On the LB/MC/IPTG plate, we found that the strain expressing NtrX^D53A^ or NtrX^D53N^ did not form visible colonies; however, the strain expressing NtrX^D53E^ formed some colonies (Fig. S1). We observed GFP-labeled *S. meliloti* cells (32) cultured in LB/MC/IPTG broth up to the logarithmic phase under a fluorescence microscope and found that more than 20% of Sm1021/p*ntrX^D53E^* cells had abnormal morphology, while Sm1021/p*ntrX* cells were normal (Fig. 1F). The growth curve determination also showed that the growth of Sm1021/p*ntrX^D53E^* in LB/MC/IPTG broth was apparently slower than that of Sm1021/p*ntrX* (Fig. 1G). Flow cytometric analysis showed that synchronized Sm1021/p*ntrX* cells were haploid and grew for 90 min in LB/MC/IPTG broth, and were mainly diploid; cells grown for 180 min reverted to haploids (Fig. 1G). Synchronized Sm1021/p*ntrX^D53E^* cells are haploids, while cells grown in LB/MC/IPTG broth for 90 minutes were mainly diploids (Fig. 1G). However, most of the cells cultured for 180 min were haploid and diploid, but some cells were polyploid (Fig. 1G). These results suggest that the phosphorylated NtrX protein is required for the regulation of cell division of *S. meliloti*.

### Transcription of cell cycle regulated genes under the regulation of NtrX in *S. meliloti*

Since NtrX is involved in controlling cell division of *S. meliloti*, does it regulate the transcription of cell cycle regulatory genes such as *ctrA, gcrA* and *dnaA?* To answer this question, we applied quantitative RT-PCR technology to analyze the transcript levels of cell cycle regulatory genes in *S. meliloti* cells. First, the synchronized Sm1021 and SmLL cells were subcultured into LB/MC broth for shaking incubation, and then total RNA was extracted from cells that grew every half hour. The qRT-PCR results showed that the transcript level of the *ntrX* gene increased first, then decreased, and reached the maximum value in the cells cultured for 90 min, showing a trend of cyclical changes, while the *ntrY* gene cyclical transcription trend was not obvious (Fig. 2A). Known cell cycle regulatory genes, such as *ctrA, gcrA* and *dnaA*, also showed a cyclical transcription trend (Fig. 2A and S2A). Compared to the wild type, in the SmLL1 mutant, transcript levels of the *ntrX* gene were lower in cells grown at different times, but the cyclical trend was unchanged, and cell cycle regulatory genes with similar down-regulation trend of *ntrX* transcription included *dnaA, ftsZ1, pleC, chpT* and *cpdR1* (Fig. 2A and S2A). Contrary to these results, *ctrA* was mainly up-regulated in SmLL1 mutants with similar trends to *gcrA, ccrM* and *ntrY* (Fig. 2A and S2A). These findings suggest that the NtrX protein may repress the transcription of genes such as *ctrA* and *gcrA* and activate the transcription of genes such as *dnaA* and *ftsZ1*.

**Fig. 2.**
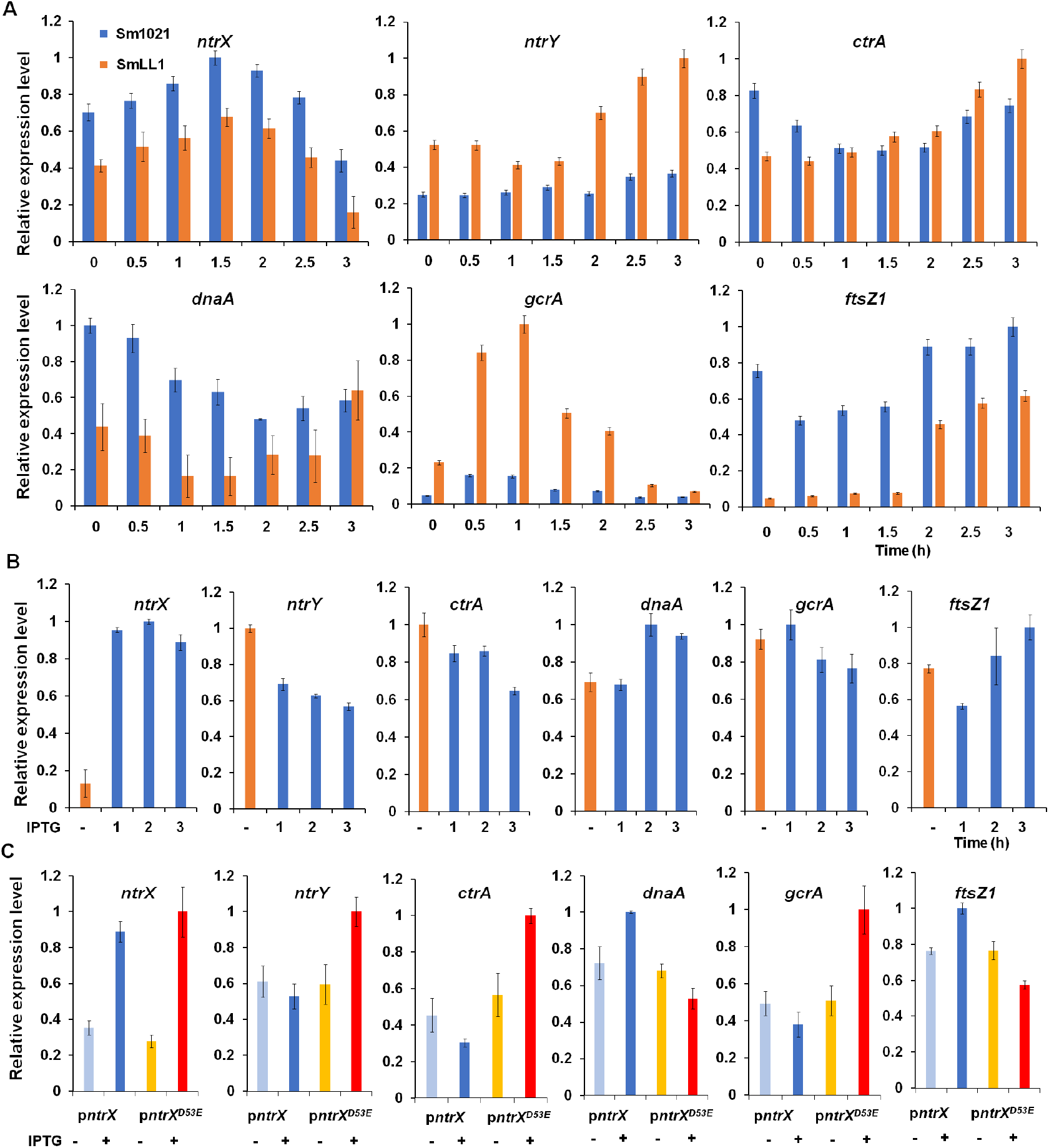
Differential expression of cell cycle regulatory genes in *ntrX* mutants. Transcript levels of cell cycle regulatory genes in *S. meliloti* cells were evaluated by qRT-PCR. Error bars, ±SD. The depletion strain was used in **B**. The genetic background was Sm1021 in **C**. 1 mM IPTG was added to the LB/MC broth in **B** and **C**.

We analyzed the transcripts of cell cycle regulatory genes in cells of the depletion strain to verify the above results. The qRT-PCR results showed that depleted cells cultured in LB/MC broth without IPTG had extremely low levels of *ntrX* transcripts, while transcripts of most cell cycle regulatory genes were available for detection (Fig. 2B and S2B). After culturing depleted cells in LB/MC broth containing IPTG for 1 h, a large number of *ntrX* gene transcripts were detected (Fig. 2B and S2B). The transcript levels of cell cycle regulatory genes were significantly altered in depleted cells cultured in IPTG broth for 2 or 3 h compared to cells cultured without IPTG broth: the transcription of *ctrA, gcrA* and *ccrM* was down-regulated; the transcription of *dnaA, ftsZ1, pleC, chpT* and *cpdR1* was up-regulated (Fig. 2B and S2B). These results further confirm that the NtrX protein represses the transcription of genes such as *ctrA* and *gcrA* and simultaneously activates the transcription of genes such as *dnaA* and *ftsZ1*.

We analyzed the transcript levels of major cell cycle regulatory genes in Sm1021/p*ntrX^D53E^* and Sm1021/p*ntrX* cells to determine whether the conserved D53 residue on NtrX is essential for transcriptional regulation. The qRT-PCR results showed that transcripts of the *ntrX* gene were significantly increased in cells cultured in LB/MC broth containing IPTG for 2 h compared to cells without IPTG; meanwhile, the transcripts of *ctrA, gcrA* and *ntrY* were significantly reduced or tended to decrease, while the transcripts of *dnaA* and *ftsZ1* genes were significantly increased (Fig. 2C). Contrary to the above results, as transcripts of the *ntrX*^D53E^ gene increased significantly under IPTG induction, transcripts of *ctrA, gcrA*, and *ntrY* also increased significantly, while transcripts of *dnaA* and *ftsZ1* genes decreased significantly (Fig. 2C). These results further confirm that the NtrX protein represses the transcription of genes such as *ctrA* and *gcrA* and activates the transcription of genes such as *dnaA* and *ftsZ1*, and that this regulation depends on the D53 residue on NtrX.

### Regulation of protein levels of CtrA and GcrA by NtrX in *S. meliloti*

To determine whether protein levels of key cell cycle regulators are affected by the *ntrX* mutation, we first expressed His-tagged NtrX, CtrA and GcrA proteins in *E. coli*. After purification by nickel columns, anti-rabbit polyclonal antibodies were prepared for immunoblotting assays (35). Sm1021-synchronized cells were subcultured in LB/MC broth for shaking incubation. Cells were collected every half hour and lysed for immunoblotting assays. The results showed a varying trend of increasing first and then decreasing NtrX protein levels and a maximum occurring in cells subcultured for 1.5 h (Fig. 3A). Unlike Sm1021, the total NtrX protein level in SmLL1 cells was apparently reduced and tended to increase gradually at different growth times (Fig. 3A). Contrary to the NtrX protein, the change trend of CtrA and GcrA protein levels in Sm1021 first decreased and then increased. The levels of these two proteins were apparently increased in SmLL1 cells compared to Sm1021 cells, (Fig. 3A). These results indicate that SmLL1 is a down-regulated mutant of *ntrX* and that NtrX protein levels are negatively correlated with CtrA and GcrA proteins.

**Fig. 3.**
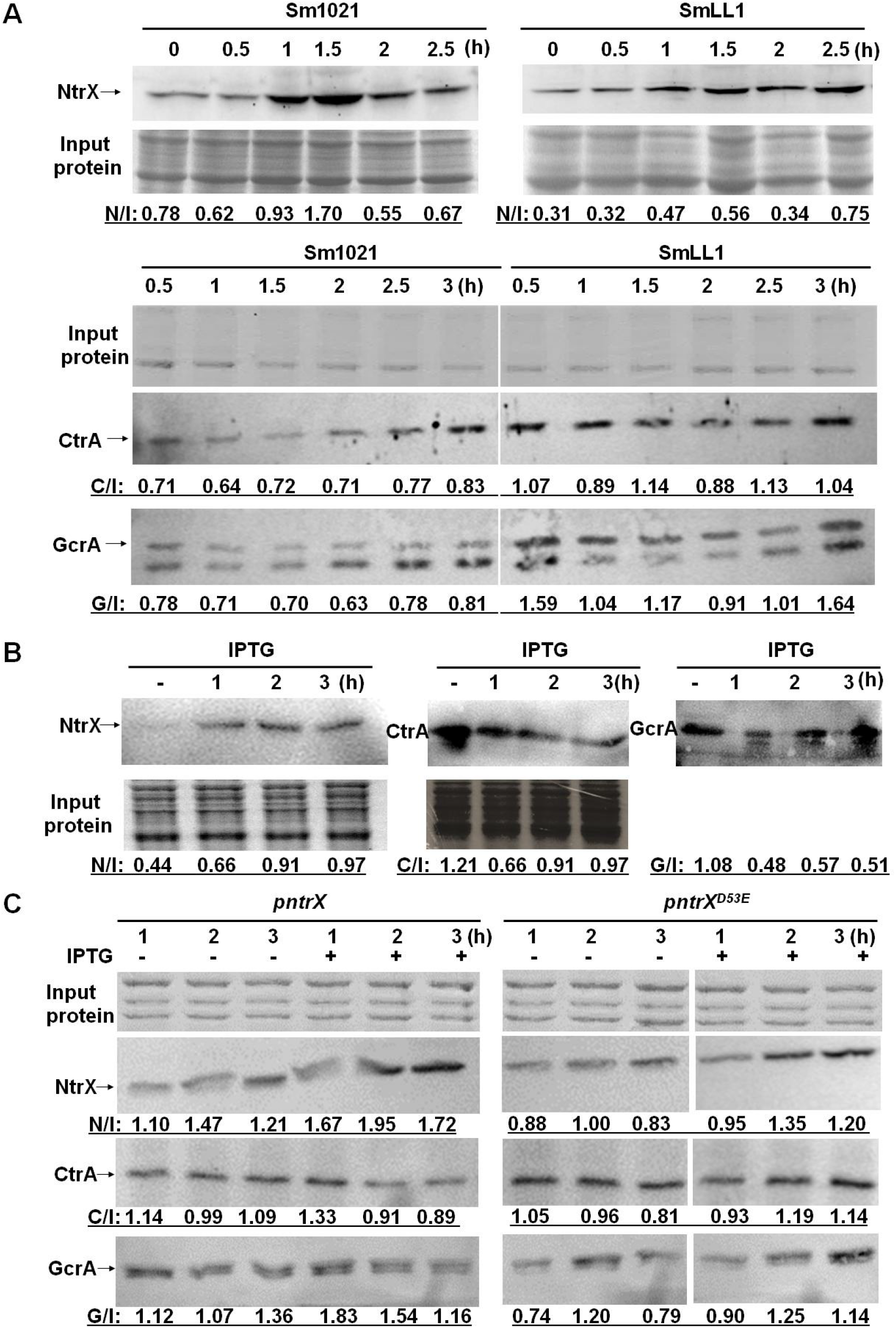
Protein levels of NtrX, CtrA and GcrA in the *ntrX* mutant as evaluated by Western blotting. N/I, C/I or G/I, the blot intensity of NtrX, CtrA or GcrA (the larger one)/the blotting intensity of the input proteins, as collected in the original data by Image J. The depletion strain was used in **B**. The genetic background was Sm1021 in **C**. 1 mM IPTG was added to the LB/MC broth in **B** and **C**.

We evaluated the NtrX protein level in cells of depleted strains grown in LB/MC broth by immunoblotting and found that cells cultured in the broth containing IPTG for 1 to 3 h expressed high levels of NtrX protein (Fig. 3B). CtrA and GcrA protein levels were highly expressed in cells cultured in LB/MC broth without IPTG, whereas the levels of these two proteins were reduced in cells cultured in broth containing IPTG for 1 to 3 h (Fig. 3B). These results also prove that NtrX protein levels are negatively correlated with CtrA and GcrA proteins.

To determine whether the D53 residue on the NtrX protein affects the protein levels of CtrA and GcrA, we performed immunoblot assays on Sm1021/p*ntrX*^D53E^ and Sm1021/p*ntrX* cells. The results showed that the protein levels of NtrX and NtrX^D53E^ increased significantly when cultured in LB/MC broth containing IPTG for 2-3 h (Fig. 3C). Under the same culture conditions, the protein levels of CtrA and GcrA were apparently reduced in Sm1021/p*ntrX* cells from the same cultured condition, while they were elevated in Sm1021/p*ntrX*^D53E^ cells (Fig. 3C). These results reaffirm that the level of NtrX protein is negatively correlated with CtrA and GcrA proteins, depending on D53 residue on the NtrX protein.

### The 53^rd^ aspartate residue as a phosphorylation site of *S. meliloti* NtrX

The homologous NtrX proteins in α-proteobacteria are composed of REC and DNA binding domains. The three-dimensional structure of the NtrX protein from *B. abortus* has been partially resolved (27–28). Using this as a template, we reconstructed the 3D structure of the NtrX protein from *S. meliloti* and found that there were 5 α-helices and 5 β-sheets connected by loops in the REC domain (Figure 4A-B). The conserved D53 is located at the end of the third β-sheet and can form a negative charge center with D9, D10 and E11 (Figure 4A-B). Mutation of D53 into A and N, but not E, may disrupt the formation of this negative charge center.

**Fig. 4.**
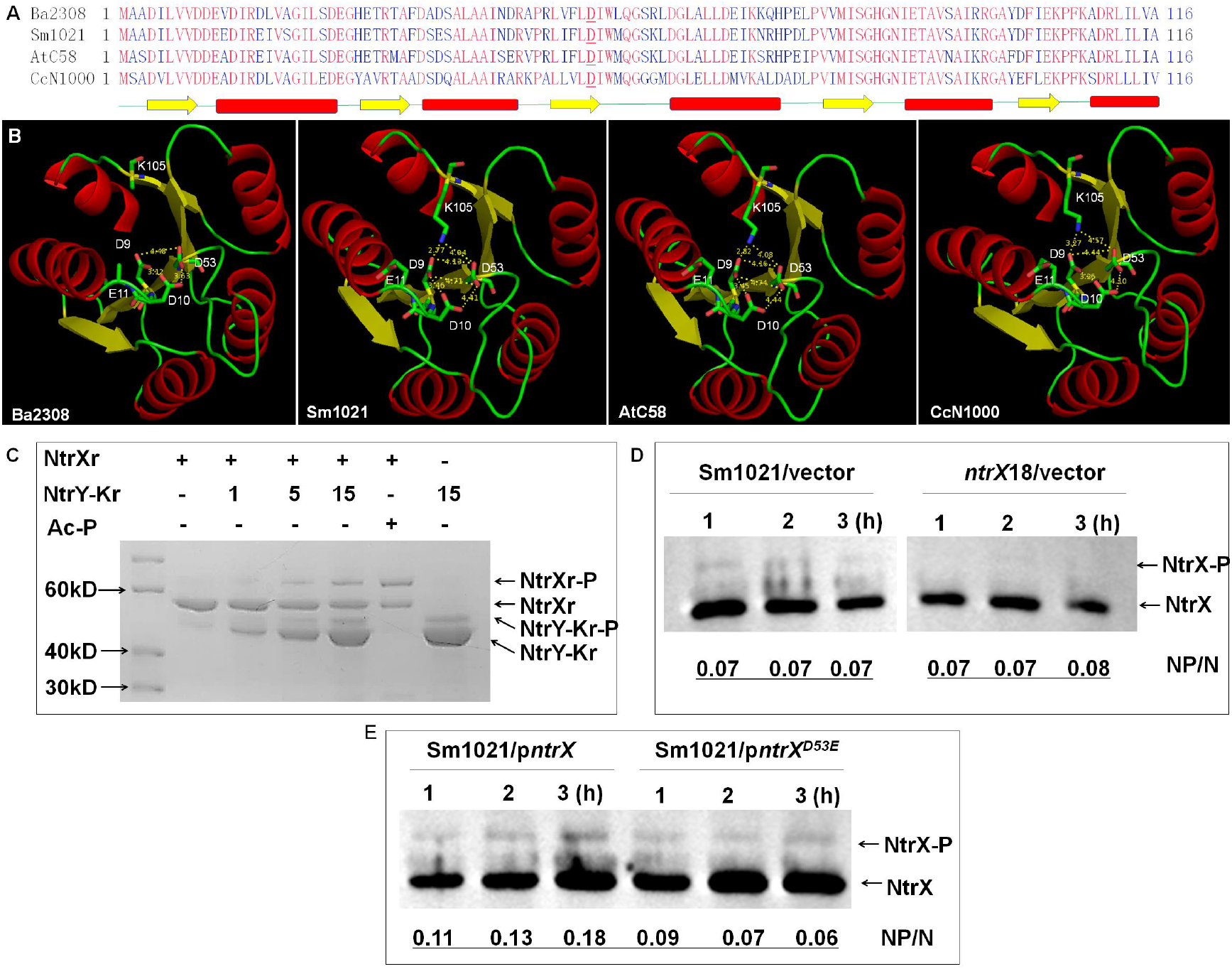
Phosphorylation of the D53 residue in the NtrX protein. (**A**) Alignment of f NtrX receiver domain from four bacterial species. The amino acid sequence of each protein was obtained from NCBI. Secondary structures of the receiver domain are shown as green lines (loops), yellow arrows (β-sheets) and red bars (α-helixes). Ba2308, *B. abortus* bv. 1 str. 2308; Sm1021, *S. meliloti* 1021; AtC58, *A. tumefaciens* C58; CcN1000, *C. crescentus* N1000. (**B**) 3D structures of the NtrX receiver domain from three bacterial species were reconstructed using *B. abortus* homolog protein (PDB:4d6y) as a template in Swiss-Model. Electrostatic interactions of the carboxyl group from the conserved 53^rd^ aspartate residues are shown as Arabic numbers via Pymol. (**C**) *In vitro* NtrX phosphorylation catalyzed by the NtrY kinase domain. NtrY-Kr, His-NtrY kinase domain fusion protein; NtrXr-P, phosphorylated NtrXr; Ac-Pi, acetyl phosphate. (**D-E**) Phosphorylated NtrX proteins from *S. meliloti* cells detected in Phos-Tag gel and by Western blotting. *S. meliloti* 1021 carrying p*ntrX* or p*ntrX*^D53E^ cultured in LB/MC broth was induced by 1mM IPTG for 1 to 3 h. ~ 1 μg of total protein was input for each sample. P/NP, intensity of phosphorylated protein/intensity of non-phosphorylated protein, as collected in the original data by Image J.

The NtrY histidine kinase can phosphorylate NtrX from *B. abortus in vitro* (27–28). We also expressed and purified the NtrY kinase domain and NtrX protein of *S. meliloti* in *E. coli* for *in vitro* phosphorylation assays. Through Phos-Tag gel detection, we found that the NtrY kinase domain can be autophosphorylated and can also phosphorylate the NtrX protein *in vitro* (Fig. 4C). After mutating D53 to E, phosphorylated NtrX protein could not be detected (data not shown). To further determine whether NtrX was phosphorylated in cells, both Phos-Tag gel and western blotting analysis were used. The results showed that the ratio of phosphorylated NtrX protein compared to non-phosphorylated protein in both Sm1021 and SmLL1 remained unchanged, though fewer NtrX proteins were detected in SmLL1 compared to SmLL1 (Fig. 4D). To further verify that the D53 residue is the phosphorylation site of the NtrX protein, we applied the same method to analyze the phosphorylated NtrX protein level of Sm1021/p*ntrX*^D53E^ cells cultured in LB/MC/IPTG broth. The results showed that the ratio of phosphorylated NtrX protein compared to unphosphorylated protein in Sm1021/p*ntrX* cells tended to increase, while the ratio in Sm1021/p*ntrX*^D53E^ cells tended to decrease (Fig. 4D). Our results suggest that NtrX can be phosphorylated in *S. meliloti* with the phosphorylation site being the D53 residue.

### Direct binding of phosphorylated NtrX protein to the promoter regions of key cell cycle regulatory genes

To determine whether the NtrX protein of *S. meliloti* directly regulates the expression of cell cycle regulatory genes, we used anti-NtrX polyclonal antibodies to perform chromatin immunoprecipitation experiments. Sequencing results showed that a total of 82 DNA fragments were specifically precipitated from Sm1021 cells, 60 of which were derived from the chromosome, while the other 22 fragments originated from the plasmids SymA (10) and SymB (12, Table S1). After careful sequence analysis of the precipitated DNA fragments, we found that the promoter regions of some cell cycle regulatory genes (such as *ctrA, dnaA* and *ftsZ1*) were enriched (Fig. 5A). Using the MEME program to scan the precipitated DNA fragments, we found that some gene promoter regions contained CAN5TG motifs, which are similar to the characteristic sequence of the *ntrY* gene promoter (CAAN3-5TTG) reported in *B. abortus* (28). Therefore, we used this sequence as a primer to scan the whole genomic DNA of *S. meliloti* 1021 by running the genome-scan dna-pattern program on the RSAT website (http://embnet.ccg.unam.mx/rsat), and a total of 2155 CAAN_1-5_TTG motifs were identified (Table S2). Fourteen of these motifs are located in the promoter regions of some cell cycle regulatory genes, such as *ctrA, danA, gcrA* and *ftsZ1*(Fig. 5B). These results suggest that the NtrX protein may recognize CAAN1-5TTG motifs in the promoter regions of some cell cycle regulatory genes to directly regulate their transcription. To verify the ChIP-Seq results, we applied quantitative PCR to evaluate the level of genomic DNA fragments from Sm1021 cells enriched with anti-NtrX polyclonal antibodies. The results showed that the promoter regions of *ctrA, dnaA, gcrA* and *ftsZ1* genes were precipitated to different degrees (Fig. 5C), indicating that the NtrX protein in Sm1021 cells can interact directly with the promoter regions of the aforementioned cell cycle regulatory genes.

**Fig. 5.**
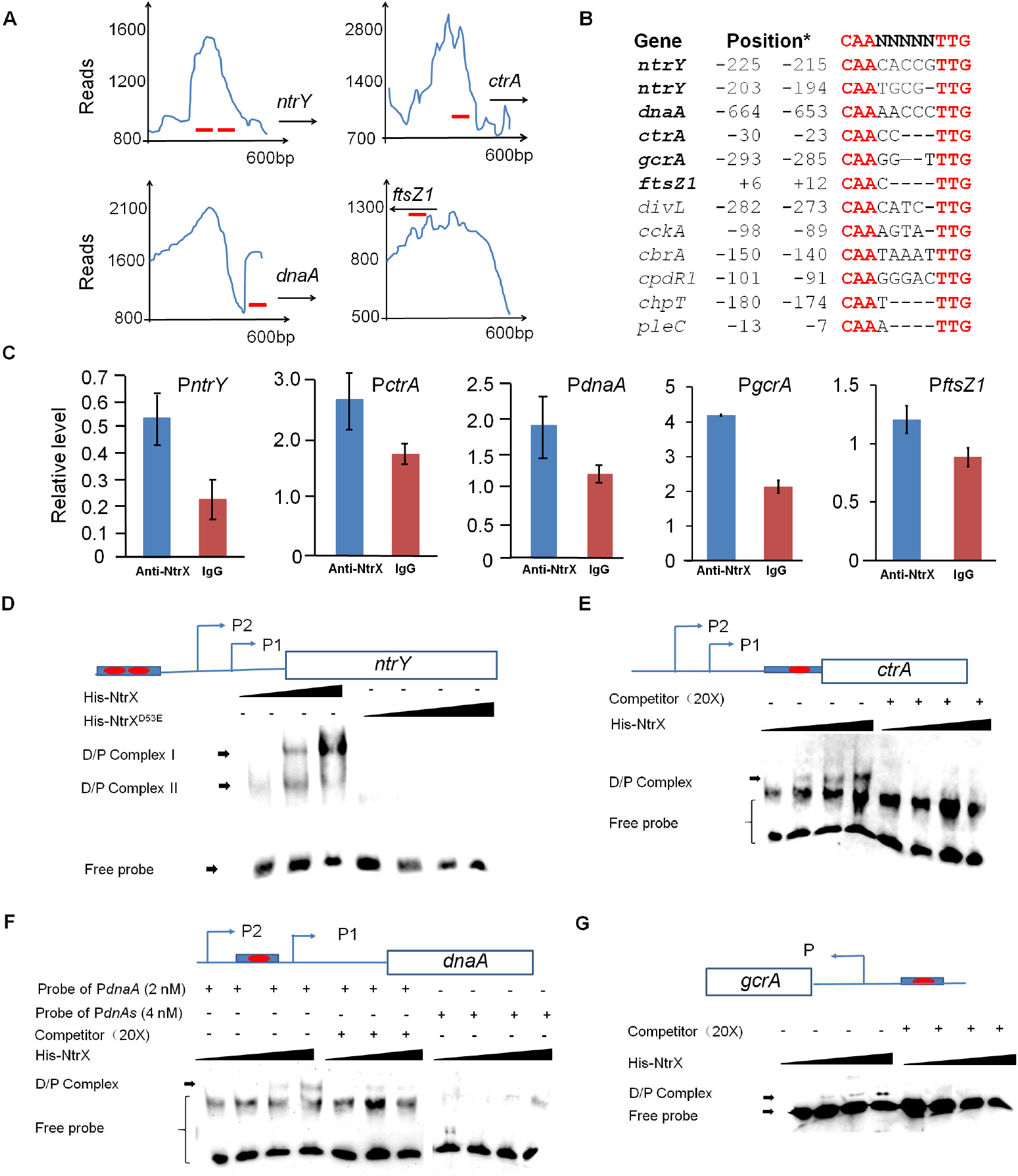
Phosphorylated NtrX binds to the promoter regions of key cell cycle regulatory genes. (**A**) Peak maps showing the promoter regions of some cell cycle genes precipitated by anti-NtrX antibodies in two independent ChIP-Seq experiments. (**B**) The conserved motif CAAN(_1-6_) TTG in the promoters of cell cycle regulatory genes. *, Predicted translation initiation as indicated by +1. (**C**) Abundance of DNA fragments specifically precipitated by anti-NtrX antibodies, as determined by qPCR. (**D-G**) The promoter regions of *ntrY* (**D**), *ctrA* (**E**), *dnaA* (**F**), and *gcrA* (**G**) containing at least one CAAN(_1-6_) TTG motif are directly bound to the phosphorylated NtrX protein *in vitro*, as based on EMSA. His-NtrX^D53E^, the His-NtrX fusion protein containing a substitution of D53 (phosphorylation site) for E in (**D**). Probe *PdnaAs* is the DNA probe *PdnaA*, where C**AA**AACCC**TT**G is replaced by C**GG**AACCC**CC**G in (**F**). D/P complex, DNA-protein complex; competitor, DNA is probed without biotin labeling. The amounts of His-NtrX proteins per probe (2 nM) are 3, 6 and 15 ng, respectively. P, putative transcriptional start sites as predicted by the BDGP program. Blue bars, probes for EMSA; red bars, putative binding sites.

In Sm1021, the promoter region of the *ntrY* gene can directly interact with the NtrX protein (Fig. 5A-C), which is consistent with the report in *B. abortus* (28). To further confirm the above findings, we synthesized a biotin-labeled *ntrY* gene promoter DNA probe (containing two **CAAN**3-5**TTG** motifs: **CAA**CACCG**TTG** and **CAA**TGCG**TTG**) for gel retardation assays. The results showed that phosphorylated NtrX specifically bound to it, forming two protein-DNA complexes (Fig. 5D). To determine whether the D53 phosphorylation site of the NtrX protein is involved in the protein-DNA binding reaction, we replaced the phosphorylated NtrX protein with the NtrX^D53E^ protein. The gel retardation results showed almost no protein-DNA complex formation (Fig. 5D), indicating that phosphorylation at position D53 is essential for the binding of NtrX to the *ntrY* promoter region. The same method was used to analyze the binding ability between the phosphorylated NtrX protein and the biotin-labeled *dnaA* promoter DNA probe (containing the **CAA**AACCC**TTG** motif) and found that they bound specifically to form a protein-DNA complex (Fig. 5F). We mutated the **CAA**AACCC**TTG** motif of the DNA probe to **CGG**AACCC**CCG**, and found that the mutant probe virtually did not bind to the phosphorylated NtrX protein (Fig. 5F), indicating that the AA or TT bases conserved in the **CAA**AACCC**TTG** motif of the *dnaA* promoter region are required for the probe binding to the phosphorylated NtrX protein. In addition, we used gel retardation assays to confirm that the phosphorylated NtrX protein can specifically bind to biotin-labeled *ctrA, gcrA* and *ftsZ1* promoter probes (each containing a **CAAN**_1-5_**TTG** motif: **CAA**CC**TTG**, **CAA**ACC**TTG** and **CAA**C**TTG**), and at least one protein-DNA complex is formed, respectively (Fig. 5E, G and S3). These results indicate that the phosphorylated NtrX protein can bind specifically to the promoter regions of *ctrA, gcrA, dnaA* and *ftsZ1 in vitro*.

### The role of *S. meliloti* NtrX proteins in *Agrobacterium tumefaciens*

The NtrX proteins from *A. tumefaciens* C58 and *S. meliloti* 1021 are highly homologous (Fig. 4A-B). To determine whether the function of NtrX protein in regulating cell division is conserved in bacteria, we introduced plasmids expressing *S. meliloti* NtrX or its substitution mutants into *A. tumefaciens* C58. By observing the growth phenotype of the transformed strain on the LB/IPTG agar plate, we found that the strain carrying p*ntrX* or p*ntrX*^D53E^ grew slower than the strain with the control vector, p*ntrX*^D53A^ or p*ntrX*^D53N^ (Fig. S4), indicating that the expression of NtrX or NtrX^D53E^ (partially retained functions) proteins can suppress cell division of *A. tumefaciens* C58, while suggesting that the NtrX protein of *A. tumefaciens* C58 is also involved in the cell cycle regulation.

Since NtrX recognizes the conserved **CAAN**_1-5_**TTG** motif, do the promoter regions of cell cycle regulatory genes from *A. tumefaciens* C58 contain these motifs? To answer this question, we used the sequence as a primer to scan the whole genomic DNA of *A. tumefaciens* C58 by running the genome-scan dna-pattern program on the RSAT website (http://embnet.ccg.unam.mx/rsat), and a total of 2562 **CAAN**_1-5_**TTG** motifs were identified. Among them, 9 motifs were located in the promoter regions of the cell cycle regulatory genes *ctrA, dnaA, gcrA, ftsZ1* and *ftsZ2*, respectively (Fig. S5). This result suggests that the NtrX protein in *A. tumefaciens* C58 may be involved in cell cycle regulation by recognizing these motifs in the gene promoter regions.

## DISCUSSION

In symbiotic nitrogen-fixing bacteria, rhizobia, the level of combined nitrogen as a signal not only regulates the expression of nitrogen-fixing genes, but also can affect cell growth and division. However, the molecular mechanism by which combined nitrogen levels regulate bacterial cell division is unclear. This work first revealed in *S. meliloti* that the phosphorylated nitrogen metabolism regulator NtrX directly regulates the transcription of cell cycle regulatory genes such as *ctrA, gcrA, dnaA* and *ftsZ1* by specifically recognizing the characteristic sequence of the promoter region (**CAAN**1-5**TTG**) and promotes cell division (Fig. 6), which provides a preliminary answer to the above question.

**Fig. 6.**
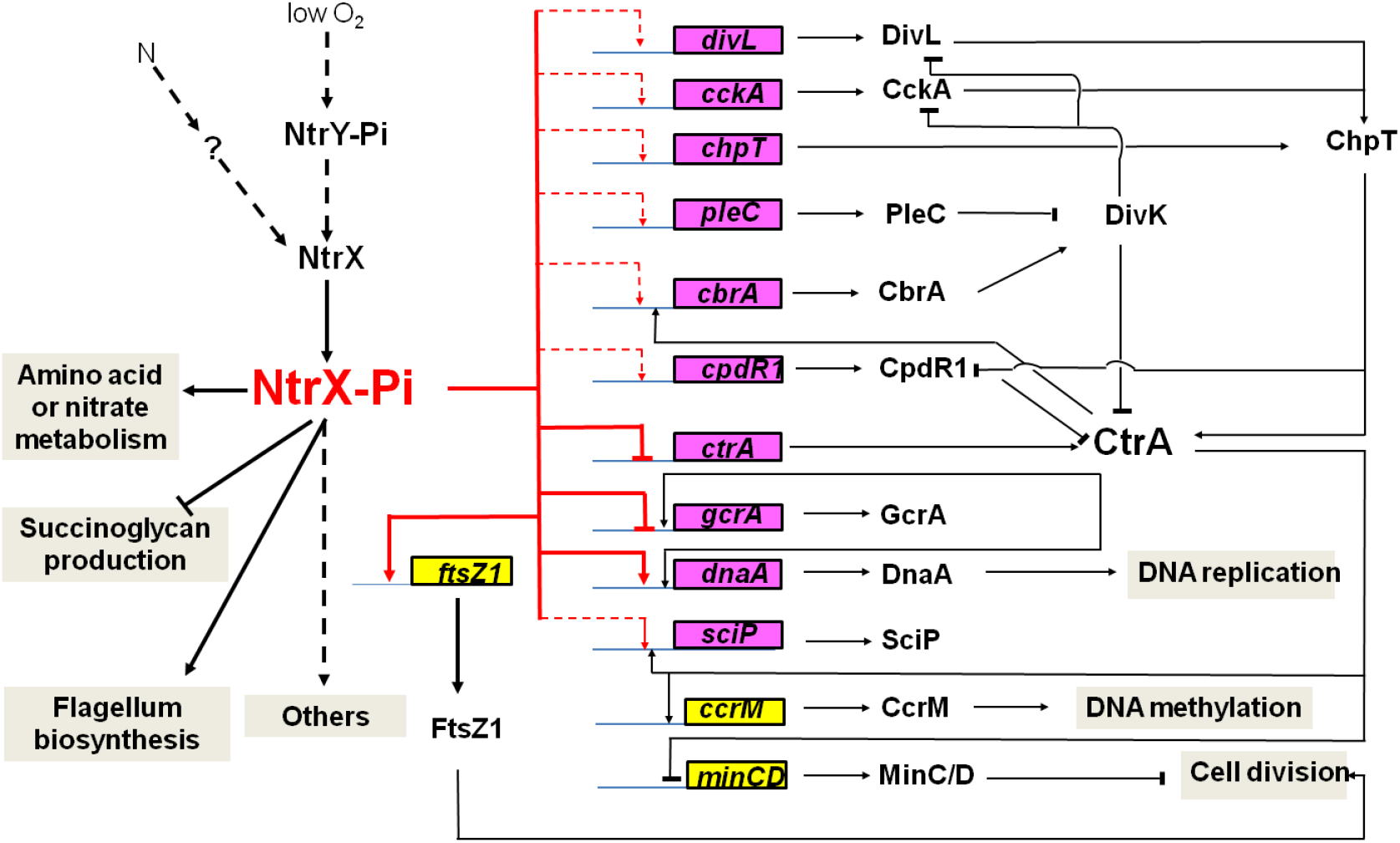
An NtrX-mediated transcriptional control system for cell cycle progression of *S. meliloti*

NtrX is a bacterial cell cycle regulator. Previous studies have suggested that NtrX is a regulator of nitrogen metabolism in bacterial cells because its mutants affect the utilization of nitrogen sources (17–20, 23, 25). Decreased utilization of nitrogen sources would inevitably lead to weakened nucleic acid and protein synthesis, which would subsequently affect the growth and proliferation of bacterial cells. This is one explanation for the effect of NtrX on bacterial cell division. In *E. chaffeensis* cells, NtrX affected the stability of CtrA protein through a post-translational mechanism (25), indicating that NtrX may act directly on the cell cycle regulatory system for regulating cell division. This work was carried out using *S. meliloti* as the study material to reveal the transcriptional control mechanism of the cell cycle regulatory genes through which NtrX protein affects cell division for the first time. This conclusion is supported by multiple experimental evidences: 1) all three groups of *ntrX* gene mutant materials are defective in bacterial growth, cell division and genomic DNA synthesis (Fig. 1); 2) the transcript levels of cell cycle regulatory genes are differentially altered in *ntrX* mutants (Fig. 2; 29); 3) the protein levels of CtrA and GcrA are altered in the mutant and they were negatively correlated with the level of NtrX protein (Fig. 3); 4) the NtrX protein is phosphorylated *in vivo* and bind directly to the promoter regions of *ctrA, gcrA, dnaA* and *ftsZ1* (Fig.4–5). The degree of cell division defects and differential gene expression may be related to the level of *ntrX* gene expression in different genetic materials and the functional status of the NtrX protein. The ability of phosphorylated NtrX in binding to DNA may be associated with the difference in the composition of the target sequence (including the recognition site).

NtrX affects cell division through transcriptional regulation of the cell cycle control system, which may also exist in other α-proteobacteria. *A. tumefaciens* and *S. meliloti* belong to the *Rhizobium* family, and their NtrX proteins are highly homologous (Fig.4A-B). We initially speculated that *S. meliloti*’s NtrX might play the same role in *A. tumefaciens* cells, but heterologous expression results showed that *S. melilot*’s NtrX or NtrX^D53E^ proteins suppressed the growth and proliferation of *A. tumefaciens* cells, while NtrX^D53A^ or NtrX^D53N^ protein did not (Fig. S4). This may reflect that the different modes of action of NtrX proteins in these two species of bacteria, which may include requirements for NtrX phosphorylation, target genes for transcriptional regulation, and even the number and location of recognition sites for the same cell cycle regulatory genes (Fig. S4). NtrX in *C. crescentus* has been reported not to affect cell growth and division during the logarithmic phase(26). By comparing the amino acid sequences of NtrX proteins, we found that the NtrX protein of *C. crescentus* is not highly homologous to that of *S. meliloti* (Fig. 4A-B), so the functions of NtrX proteins may differ between the two species of bacteria. We attempted to express the NtrX protein of *S. meliloti* in *C. crescentus*, but failed to screen transformed cells, whereas cells expressing the NtrX^D53^ substitution protein were successfully screened (data not shown), suggesting that the endogenous NtrX protein of *C. crescentus* may be an inactivated mutant protein whose function may be replaced by other proteins. Whether the NtrX protein of *B. abortus* affects cell division requires further study.

The NtrY kinase derived from *S. meliloti* and *C. crescentus* can phosphorylate NtrX protein *in vitro* (23; Fig.4-1), but there is no experimental evidence to demonstrate the phosphorylation of NtrX by NtrY in the cell. Our previous study showed no correlation between the genetic phenotypes of *ntrY* and *ntrX* mutants in *S. meliloti* 1021: the *ntrY* deletion mutant did not affect the growth and proliferation of bacterial cells and inoculated hosts of alfalfa were not defective in symbiotic nitrogen fixation, whereas the SmLL1 mutant grew slowly in LB/MC and inoculated host plants formed inefficient nitrogen-fixing nodules (29). Therefore, the upstream kinase of NtrX in the cell may not be NtrY. This inference is consistent with the repression of *ntrY* gene expression by NtrX in *S. meliloti* (29; Fig. 2A). Furthermore, the study results from *B. abortus* indicate that the expression of *ntrY* gene is induced by micro-oxygen and is involved in micro-oxygen signal transduction (23). The primary function of NtrX in bacteria is considered to regulate nitrogen metabolism. The nitrogen limitation signal transduction in bacteria is mainly mediated by the NtrB/NtrC system (33), so it cannot be ruled out that the NtrB/NtrC system can regulate the expression of *ntrX* under nitrogen lacking conditions. Under nitrogen rich conditions, the activity of NtrX may be regulated by an unknown kinase, which may be able to sense the level of ammonia nitrogen.

Finally, *S. meliloti* can infect alfalfa and induce the formation of nitrogen-fixing nodules. On the root surface of the plant, rhizobia cells grow and divide at the top of the extended infection thread and in the nodule infection zone. Whether these division events are regulated by NtrX requires further study.

## MATERIALS AND METHODS

### Strains and culture medium

*Escherichia coli* DH5α and BL21 were cultured in LB medium at 37 °C. *S. meliloti* (including Sm1021, SmLL1 and derivatives) (29) were cultured in LB/MC medium at 28 °C. MOPS-GS broth was utilized for the cell synchronization of *S. meliloti* (34). *A. tumefaciens* C58 was cultured in LB medium at 28 °C. The following antibiotics were added to the medium: kanamycin, 50 μg/ml; gentamicin, 10 μg/ml; chloramphenicol, 30 μg/ml; neomycin, 200 μg/ml; streptomycin, 200 μg/ml; tetracycline, 10 μg/ml.

### Recombinant plasmid construction

Primers P*ntrX*1 and P*ntrX*2 bearing *Hind*III and *Xba*I digestion sites were used to amplify the *S. meliloti ntrX* gene (Table S3). The *ntrX* gene fragment was amplified using overlapping PCR primers NMF and NMR with substitution of aspartate for glutamate, asparagine or alanine (Table S3). Overlapping PCR was performed as described by Wang(29) in 2013. The PCR products were digested with *Hind*III and *Xba*I (Thermo) and ligated with digested pSRK-Gm (31) to obtain the recombinant plasmids p*ntrX*, p*ntrX^D53E^*, p*ntrX*^D53A^ and p*ntrX*^D53N^. Upon introducing p*ntrX* into SmLL1, the depletion strain was screened on LB/MC agar plates containing 1 mM IPTG. Primers for P*ntrY*k1, P*ntrY*k2, p*ntrX*1^D53E^, p*ntrX*2^D53E^, P*ctrA*1, P*ctrA*2, P*gcrA*1 and P*gcrA*2 were used to amplify the NtrY kinase domain, *ntrX*^D53E^, *ctrA* and *gcrA*, respectively (Table S3). The PCR fragments were digested with the appropriate restriction enzymes and ligated into pET28b to harvest p*ntrY*k, p*ntrX*^D53E^, p*ctrA* and p*gcrA* for recombinant protein purification.

### Bacterial cell synchronization

De Nisco’s method was used for bacterial cell synchronization (34). A sample of *S. meliloti* cells was selected from an agar plate, placed in 5 ml LB/MC broth and grown overnight. A 100-μl aliquot of the bacterial culture was transferred to 100 ml LB/MC broth and incubated overnight until OD_600_ = 0.1-0.15; the culture was centrifuged (6,500 rpm, 5 min, 4°C) and the cell pellet was washed twice with sterilized 0.85% NaCl solution. Cells were resuspended in MOPS-GS synchronization medium and grown at 28°C for 270 min. After centrifugation, cells were washed twice with sterilized 0.85% NaCl solution, resuspended in LB/MC broth, and cultured at 28°C.

### RNA extraction and purification

20 ml of bacterial cultures were collected by centrifugation (6,000 rpm, 5 min, 4°C) and the cells were washed twice with DEPC-treated water. RNA extraction was performed using 1 ml of Trizol (Life Technology) for RNA extraction, and its purification was performed as described by Wang (29).

### qRT-PCR and qPCR

RNA reverse transcription was performed using the PrimeScript RT reagent Kit with gDNA Eraser (TaKaRa). The qPCR reaction system included the following: SYBR^®^ Green Real-time PCR Master Mix, 4.75 μl; cDNA or DNA, 0.25 μl; 10 pmol/μl primers, 0.5 μl; ddH_2_O, 4.5 μl. The reaction procedure is as follows: 95°C, 5 min; 95°C, 30 s; 55°C, 30 s; 72°C, 1 kb/min. The selected reference gene was Smc00128. The 2-^ΔΔ^CT method was applied to the analysis of gene expression levels. All primers are listed in Table S3.

### Chromatin immunoprecipitation (ChIP)

ChIP was performed as described by Pini (4). Anti-NtrX antibodies prepared by Wenyuange (Shanghai) were used for ChIP assays (35).

### Flow cytometry

Four milliliters of fresh bacterial cultures were centrifuged (6,000 rpm, 5 min, 4°C) and the cell pellets were washed twice with a 0.85% NaCl solution (stored at 4°C). De Nisco’s flow cytometry protocol of was used (34). Each sample was assessed using a MoFlo XDF (Beckman Coulter) flow cytometer, and the results were analyzed using Summit 5.1 software (Beckman Coulter).

### EMSA (electrophoretic mobility shift assay)

EMSA was performed as described by Zeng (35). The LightShift™ Chemiluminescent EMSA Kit (Thermo Fisher) was applied in the assays. Probes for *ntrY, ctrA, dnaA, gcrA* and *ftsZ1* promoter DNA labeled with biotin were synthesized (Invitrogen); the probes are listed in Table S3. Phosphorylated NtrXr proteins (from Ni^2+^ column purification) used for these assays were prepared. 6 μg of purified protein was mixed with 2 mM of acetyl phosphate (Sigma) in 100 μl of phosphorylation buffer (50 mM Tris-HCl pH 7.6, 50 mM KCl, 20 mM MgCl_2_), and incubated for 1 h at 28°C. Acetyl phosphate was removed using an ultra-filtration tube (Millipore) resolved in 100 μl of phosphorylation buffer.

### NtrX phosphorylation assay

1 mg of His-NtrX fusion protein (NtrXr) and 1 mg of His-NtrY kinase domain fusion protein (NtrY-Kr) purified through a Ni^2+^ column were used for *in vitro* NtrX phosphorylation assays. 2 mM of acetyl phosphate (Sigma) was mixed with 300 μg of NtrY-Kr in 1 ml of phosphorylation buffer and then incubated for 1 h at room temperature. Acetyl phosphate was removed using an ultra-filtration tube (Millipore) and resolved in 1 ml of phosphorylation buffer. 1, 3 and 10 μg of phosphorylated NtrY-Kr protein were added to 200 μl of phosphorylation buffer having 10 μg of NtrXr, and incubated overnight at 28°C. Samples were separated by a Phos-Tag gel (Mu Biotechnology, Guangzhou). Phosphorylated NtrX levels from *S. meliloti* cells were separated in a Phos-Tag gel. *S. meliloti* 1021 carrying p*ntrX* or p*ntrX*^D53E^ were cultured in LB/MC broth induced by 1 mM IPTG for 1 to 3 h. ~1 μg of total protein was input for each sample. The total protein was transferred to PVDF, and then detected by anti-NtrX antibodies from Western blotting assays (13).

### Western blotting

Western blotting was performed as described by Tang (36). Proteins were detected using an ECL fluorescence colorimetric kit (Tiangen) and visualized using a Bio-Rad Gel Doc XR. Anti-NtrX antibodies were prepared at Wenyuange, Shanghai, China (35); Anti-CtrA and anti-GcrA antibodies were prepared by Hua’an Biotech, Hangzhou, China.

### Microscopy

A 5-μl aliquot of fresh *S. meliloti* culture (OD_600_ = 0.15) was placed on a glass slide and covered with a cover glass. The slide was slightly baked near the edge of the flame of an alcohol lamp for a few seconds. Cells expressing the pHC60 plasmid were observed in GFP mode (32), and images were acquired using a CCD camera Axiocam 506 color (Zeiss). Exposure times were set to 10 ms and 1,000 ms in order to capture bacterial morphology and motility, respectively. Images were analyzed with ZEN 2012 software (Zeiss).

### ChIP sequencing

ChIP-Seq was performed by Bohao Biotech, Shanghai. The original data were evaluated by Bohao Biotech based on the standard protocol. Genome mapping was then performed using bowtie2 (version: 2-2.0.5) for local alignment of clean reads (37). The original genomic data of *S. meliloti* 1021 was obtained from NCBI. IGV software was employed to assess specific enrichment data based on ChIP-Seq results, and screenshots of peak maps were generated (38).

### Bioinformatic analysis

Genome-wide CAAN_1-6_TTG *cis*-acting elements of bacteria were evaluated using the genome-scan dna pattern program on the RAST online server (http://embnet.ccg.unam.mx/rsat) (39). The BDGP program was applied for promoter prediction (http://www.fruitfly.org/seq_tools/promoter.html)

The NtrX protein 3D structure was reconstructed in Swiss-Model using the 4d6y template (the 3D structure of the NtrX receiver domain from *Brucella*) in PDB (40). The 3D structure of the NtrX receiver domains was analyzed by Pymol (Delano Scientific).

## Supporting information

MS for biorxiv

MS for biorxiv

MS for biorxiv

MS for biorxiv

## ACKNOWLEDGMENTS

This research was supported by the Natural Science Foundation of China (31570241 to L.L).

## AUTHOR CONTRIBUTIONS

L. L. designed research; S. X., F. A., X. Y., L. H., S. Z., L. Z., W. Z., and N. L. performed research; S.X., F.A., J. Y., L. Y. and L. L. analyzed data; L. L. and K. O. wrote the paper.

## SUPPORTING INFORMATION

**Table S1** DNA fragments of Sm1021 cells precipitated by anti-NtrX antibodies

**Table S2** NtrX binding sites predicted by RSAT

**Table S3** DNA oligonucleotides used in this study

**Fig. S1.** Colony formation of Sm1021 expressing NtrX substituted with D53 on LB/MC/IPTG agar plates.

**Fig. S2.** Differential expression of cell cycle regulatory genes in *ntrX* mutants.

Transcript levels of cell cycle regulatory genes in *S. meliloti* cells were evaluated by qRT-PCR. Error bars, ±SD. The depletion strain was used in **B**. 1 mM IPTG was added to the LB/MC broth in **B**.

**Fig. S3.** The promoter regions of *ftsZ1* containing one CAAN(_1-6_)TTG motif bind directly to the phosphorylated NtrX protein *in vitro*.

D/P complex, DNA-protein complex; competitor, DNA is probed without biotin labeling. The amounts of His-NtrX proteins per probe (2 nM) are 3, 6 and 15 ng, respectively. P, the putative transcriptional start site was predicted by the BDGP program. Blue bars, probes for EMSA; red bars, putative binding sites.

**Fig. S4.** Growth phenotypes of *A. tumefaciens* expressing *S. meliloti* NtrX substitutions.

**Fig. S5.** Distribution of CAAN(_1-5_)TTG motifs in three α-proteobacterial species by scanning genomic DNA in RSAT.

These sites can be recognized by NtrX to affect gene transcription. *, the predicted translation start is indicated by +1.

