## Supplementary material for "Direct regulation of cell cycle regulatory gene expression by NtrX to promote *Sinorhizobium meliloti* cell division": MS for biorxiv

**SUPPORTING INFORMATION**

**Table S1** DNA fragments of Sm1021 cells precipitated by anti-NtrX antibodies

**Table S2** NtrX binding sites predicted by RSAT

**Table S3** DNA oligonucleotides used in this study

**Fig. S1.** Colony formation of Sm1021 expressing NtrX substituted with D53 on LB/MC/IPTG agar plates.

**Fig. S2.** Differential expression of cell cycle regulatory genes in *ntrX* mutants.

Transcript levels of cell cycle regulatory genes in *S. meliloti* cells were evaluated by qRT-PCR. Error bars, ±SD. The depletion strain was used in **B**. 1 mM IPTG was added to the LB/MC broth in **B**.

**Fig. S3.** The promoter regions of *ftsZ1* containing one CAAN$_{(1-6)}$TTG motif bind directly to the phosphorylated NtrX protein *in vitro*.

D/P complex, DNA-protein complex; competitor, DNA is probed without biotin labeling. The amounts of His-NtrX proteins per probe (2 nM) are 3, 6 and 15 ng, respectively. P, the putative transcriptional start site was predicted by the BDGP program. Blue bars, probes for EMSA; red bars, putative binding sites.

**Fig. S4.** Growth phenotypes of *A. tumefaciens* expressing *S. meliloti* NtrX substitutions.

**Fig. S5.** Distribution of CAAN$_{(1-5)}$TTG motifs in three α-proteobacterial species by scanning genomic DNA in RSAT.

These sites can be recognized by NtrX to affect gene transcription. *, the predicted translation start is indicated by +1.

Fig. S1.

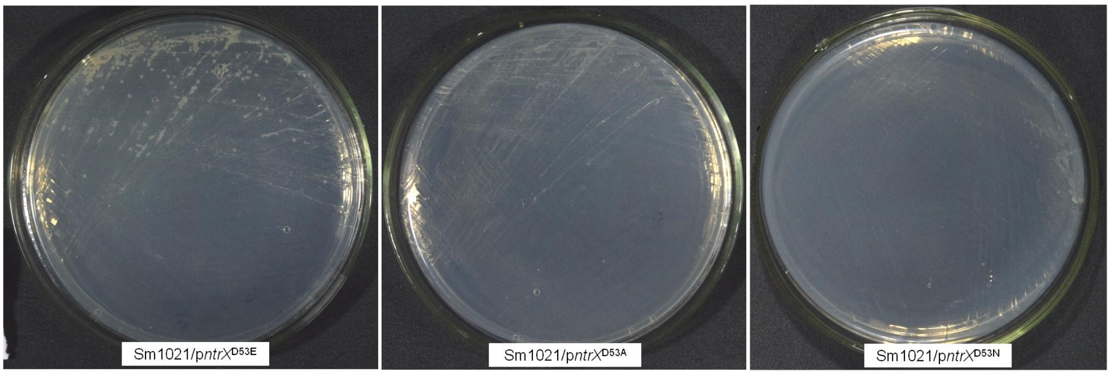

Fig. S2.

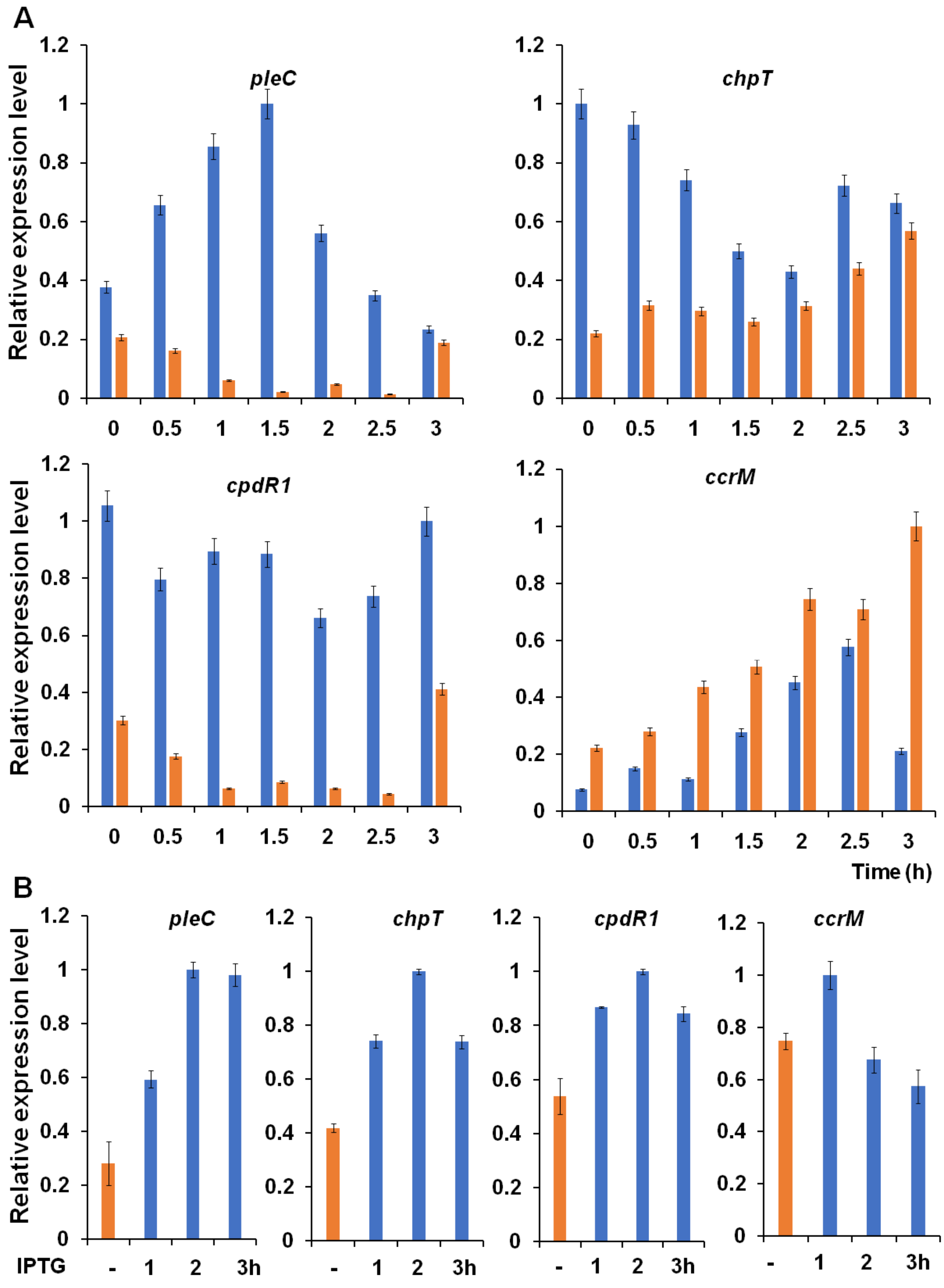

A
Relative expression level
pleC
chpT
cpdR1
ccrM
1.2
1
0.8
0.6
0.4
0.2
0
0
0.5
1
1.5
2
2.5
3
Time (h)
B
IPTG
-
1
2
3h


Fig. S3.

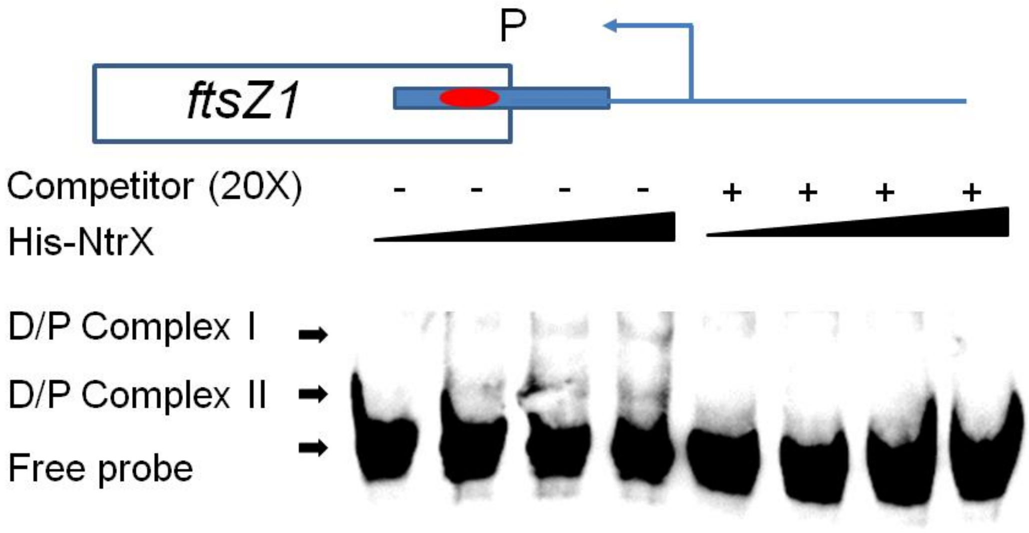
P
ftsZ1
Competitor (20X)
- - - - + + + +
His-NtrX
D/P Complex I
D/P Complex II
Free probe

Fig. S4.

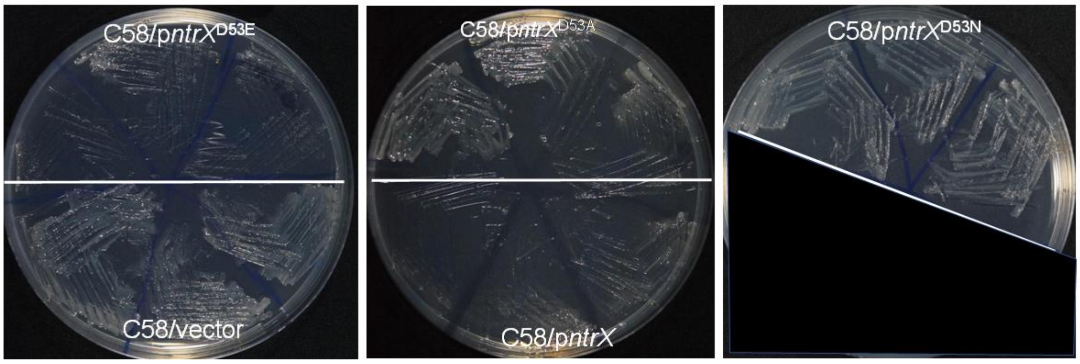
C58/p*ntrX*$^{D53E}$
C58/vector
C58/p*ntrX*$^{D53A}$
C58/p*ntrX*
C58/p*ntrX*$^{D53N}$

Fig. S5.

| Species | Gene | Position* | | CAANNNNNTTG |
|---|---|---|---|---|
| *C. crescentus* | *divJ* | -184 | -178 | CAAC----TTG |
| | *ccrM* | -123 | -113 | CAAAATAATTG |
| | *gcrA* | -182 | -173 | CAAGGTC-TTG |
| | | -333 | -323 | CAAGCGCTTTG |
| *B. abortus* | *divK* | -220 | -211 | CAACCC--TTG |
| | *dnaA* | -266 | -259 | CAAGG---TTG |
|  | *gcrA* | -208 | -202 | CAAG----TTG |
|  | *rcdA* | -225 | -219 | CAAT----TTG |
| *A. tumefaciens* | *ctrA* | -148 | -141 | CAACT---TTG |
| | | -192 | -185 | CAATT---TTG |
|  | *dnaA* | -19 | -13 | CAAT----TTG |
| | *gcrA* | -388 | -378 | CAATAGGCTTG |
| | | -239 | -230 | CAAGGTG-TTG |
|  |  | -207 | -201 | CAAG----TTG |
|  | *ftsZ1* | -63 | -57 | CAAG----TTG |
| | | -375 | -365 | CAAACGTGTTG |
| | *ftsZ2* | -139 | -130 | CAAAGGC-TTG |
